# interpolatedXY: a two-step strategy to normalise DNA methylation microarray data avoiding sex bias

**DOI:** 10.1101/2021.09.30.462546

**Authors:** Yucheng Wang, Tyler J. Gorrie-Stone, Olivia A. Grant, Alexandria D. Andrayas, Xiaojun Zhai, Klaus D. McDonald-Maier, Leonard C. Schalkwyk

## Abstract

**Motivation:** Data normalization is an essential step to reduce technical variation within and between arrays. Due to the different karyotypes and the effects of X chromosome inactivation, females and males exhibit distinct methylation patterns on sex chromosomes, thus it poses a significant challenge to normalise sex chromosome data without introducing bias. Currently, existing methods do not provide unbiased solutions to normalise sex chromosome data, usually, they just process autosomal and sex chromosomes indiscriminately.

**Results:** Here, we demonstrate that ignoring this sex difference will lead to introducing artificial sex bias, especially for thousands of autosomal CpGs. We present a novel two-step strategy (interpolatedXY) to address this issue, which is applicable to all quantile-based normalisation methods. By this new strategy, the autosomal CpGs are first normalised independently by conventional methods, such as funnorm or dasen; then the corrected methylation values of sex chromosome linked CpGs are estimated as the weighted average of their nearest neighbours on autosomes. The proposed two-step strategy can also be applied to other non-quantile-based normalisation methods, as well as other array-based data types. Moreover, we propose a useful concept: the sex explained fraction of variance, to quantitatively measure the normalisation effect.

**Availability:** The proposed methods are available by calling the function *‘adjustedDasen’* or *‘adjustedFunnorm’* in the latest wateRmelon package (https://github.com/schalkwyk/wateRmelon), with methods compatible with all the major workflows, including minfi.

**Supplementary information:** Supplementary data are available at...

## 1 Introduction

DNA methylation microarrays, such as Infinium HumanMethylation450 BeadChip (Bibikova *et al.*, 2011) and Infinium MethylationEPIC BeadChip (Moran *et al.*, 2016), provide cost-effective and high-throughput measurements of the methylation status over half a million CpG sites across the genome will continue to be the first choice by most DNA methylation related large cohort studies in the near future. Although whole genome bisulfite sequencing (WGBS) is recognized as the gold standard to measure the methylation patterns across the human genome, the high costs and technical complexity still pose significant challenges that prevent application to large-scale samples (Villicaña and Bell, 2021). Data normalisation is an important prerequisite step to reduce unwanted technical variation. Currently, several normalisation methods are available for DNA methylation microarray samples. Among them, peak-based correction (PBC) (Dedeurwaerder *et al.*, 2011), Beta MIxture Quantile normalization (BMIQ) (Teschendorff *et al.*, 2013) and noob (Triche *et al.*, 2013) are all within-array normalization methods however they do not reduce between-array variation. By contrast, dasen (Pidsley *et al.*, 2013) and funnorm (Fortin *et al.*, 2014) are the two most widely used between-array normalisation methods, which were reported to be able of effectively reducing the variation between samples. Dasen in the wateRmelon package utilises quantile normalisation to normalise methylated and unmethylated intensities separately, and also addresses Type I and Type II probes separately. Prior to the normalisation steps, there are linear regression procedures in dasen to reduce the density distribution difference between Type I and Type II probes (Pidsley *et al.*, 2013). The functional normalisation employed by funnorm is also an extension to quantile normalisation that removes variation explained by a set of selected covariates. In funnorm, the covariates are set as the first two principal components of the control probes, and linear regression is used to determine the proportion of variation explained by the covariates (Fortin *et al.*, 2014).

Females have two copies of the X chromosome, while males have one X chromosome and one Y chromosome. To compensate for the different dosages of the X chromosome genes, one X chromosome in female cells is randomly subjected to inactivation in each cell lineage, with most parts of the inactive X being highly methylated (LYON, 1961; Sharp *et al.*, 2011; Cotton *et al.*, 2015). As a result of this, the mean methylation values of the X chromosomes between sexes are very different (McCarthy *et al.*, 2014; Wang *et al.*, 2021; Grant *et al.*, 2021). The distinct methylation patterns of sex chromosomes between females and males raise a great challenge to unbiasedly normalise sex chromosome data. The existing between-array normalisation methods do not provide good solutions for normalising sex chromosome data. For example, dasen ignores this issue and normalises autosomes and sex chromosomes together, while funnorm is designed to normalise male samples and female samples separately for X chromosomes and Y chromosomes. Some DNA methylation related studies simply remove those probes mapped to the X and Y chromosomes prior to the normalisation step and do not include them in the downstream analysis. All these strategies come with their own drawbacks, either through losing some potentially interesting and biologically relevant signals from sex chromosomes or by introducing systematic technical differences between sexes.

Here we first demonstrate that the existing normalisation methods used to handle probes mapped on the X and Y chromosomes lead to introducing artificial sex bias into the normalised data. Then, we present a novel two-step strategy, which is designed to unbiasedly normalise both autosome data and sex chromosome data, is applicable to all quantile-based normalisation methods.

## 2 Materials and methods

### 2.1 Datasets

Two main datasets were used in this study. The first dataset includes 1195 individuals from the Understanding Society: UK Household Longitudinal Survey (UKHLS). Details about this UKHLS dataset are described by Gorrie-Stone et al. (Gorrie-Stone *et al.*, 2019). In brief, DNA methylation levels in whole blood within 489 male and 686 female healthy individuals were measured by EPIC array. The UKHLS dataset is available under request from the European Genome-phenome Archive under accession EGAS00001002836 (https://www.ebi.ac.uk/ega/home). Since funnorm was developed and tested on 450k array samples, in this study we produce subsets from GSE142512 (Johnson *et al.*, 2020) to evaluate funnorm. GSE142512 includes 87 individuals with type 1 diabetes (T1D) and 87 individuals without T1D. The peripheral blood samples were collected from the subjects between 1 and 5 time points, with DNA methylation levels measured by either 450K or EPIC array, further details were documented by Johnson et al. (Johnson *et al.*, 2020). We randomly selected 16 450k samples (12 males and 4 females) from GSE142512 as the dataset one which is used to evaluate the performance of funnorm on small size dataset, and randomly selected 48 450k samples (23 males and 25 females) as dataset two to test funnorm’s performance on relatively larger size dataset. For reproducibility, the sample IDs in the two subset datasets are listed in Supplementary Table 1. GSE142512 is publicly available from Gene Expression Omnibus (https://www.ncbi.nlm.nih.gov/geo/).

### 2.2 DNA methylation data process

The DNA methylation raw data (IDAT files) were read into R by either using *iadd2* function in bigmelon or *read.metharray.exp* function in minfi. The methylation level of any given CpG locus is measured by its beta value which is defined as: *β* = (*M*) / (*M* + *U* + 100), where *M* is methylated intensity and U is unmethylated intensity for a given CpG loci. Basic quality control steps were performed to identify outliers, as recommended by Gorrie-Stone et al. (Gorrie-Stone *et al.*, 2019). Further, the reported sexes of samples were checked against the predicted sexes from DNA methylation data by using the *estimateSex* function in watermelon package (Pidsley *et al.*, 2013), which predicts sex by comparing the methylation levels on sex chromosomes (Wang *et al.*, 2021). The original dasen normalisation is performed by calling the *dasen* function with default settings in the watermelon package, the original funnorm normalisation is performed by calling the *preprocessFunnorm* with default settings in the minfi package (Fortin *et al.*, 2016), which is actually applies *noob* method (Triche *et al.*, 2013) as a first step for background correction and then perform the functional normalisation.

All analyses were performed using R 3.6.0 under Linux environment.

### 2.3 A two-step strategy to unbiasedly normalise DNA methylation samples

The explicit procedures of the proposed new strategy to unbiasedly normalise both autosomal CpGs and sex chromosome linked CpGs are as follows:

1. Step one: normalise the autosomal CpGs by one of the conventional normalisation methods, such as funnorm or dasen. It should be noted, the probes mapped to sex chromosomes should not be included in this step to avoid potential influence.
2. Step two: infer the corrected values of sex chromosome linked CpGs by looking for their nearest neighbors on autosomes, this is achieved by linear interpolation, here is the very efficient implementation:

a. Sort the corrected values of autosomal CpGs and build a function *F* which reflects correspondence of the rank of a CpG to its corrected value: *Corrected_value_i_* = *F* (*rank_i_*).
b. Sort and get the ranks of autosomal CpGs based on their raw values.
c. Estimate the ranks of sex chromosome linked CpGs by linear interpolation on the rank distribution from the procedure b.
d. Put the inferred ranks of sex chromosome linked CpGs into the function *F* to get their final corrected values.

The above steps are ideally performed on raw signal intensities (M and U) and on each probe type (IGrn, IRed and II in funnorm, I and II in dasen) individually. After that, the normalised intensities can be converted into beta values as: β = (M) / (M + U + 100). We name this strategy as interpolatedXY. When dasen is used to normalise autosomal CpGs in the first step, we call this new normalisation method as “interpolatedXY adjusted dasen”. Similarly, “interpolatedXY adjusted funnorm” refers to another new normalisation method in which the functional normalisation is applied in the first step.

### 2.4 Performance evaluation for the interpolation approach

The proposed new approach infers the corrected values of sex chromosome linked CpGs by linear interpolation on autosomal CpGs. To investigate whether the inferred data is accurate and reliable, we need a gold standard to evaluate the estimation accuracy. Females and males have very different methylation patterns on sex chromosomes, that is the main reason that we avoid normalising female samples and male samples together, with autosomes and sex chromosomes treated indiscriminately. However, when the targeted dataset includes only unisexual samples (only females or only males), then the sex chromosomes should be normalised together with other autosomes.

Inspired by this, we designed single sex groups: one that includes only female samples and the second that consists of only male samples. Firstly, the two groups are both normalised by conventional methods (e.g. dasen and funnorm) with the sex chromosomes being treated as general autosomes, thus the corrected values of those sex chromosome linked CpGs could serve as the golden references (i.e. expected values). Secondly, by our proposed interpolation approach, we infer the corrected values of sex chromosome linked CpGs by interpolating on the normalised values of the autosomal CpGs. Lastly, the interpolated values are compared with their corresponding reference values. Root mean squared error (RMSE), which is sensitive to outliers, is used here to measure the deviations from the inferred values to their expected values:

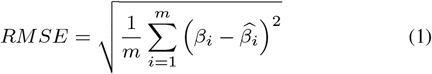

where *β_i_* is the methylation beta value of the *i^th^* CpG, 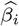 represents the expected methylation beta value of the *i^th^* CpG, *m* represents the total number of CpGs studied.

### 2.5 Evaluation of the technical sex biases

The original dasen performs quantile normalisation with autosomal CpGs and sex chromosome CpGs processed together even when the dataset to be normalised is composed of both females and males. To investigate whether such an approach would introduce artificial sex biases, we compared the normalisation results of the UKHLS dataset generated by the original dasen and the interpolatedXY adjusted dasen.

The human methylome is not constant but responsive to many internal and external factors, such as genetic backgrounds and environmental factors (Van Dongen *et al.*, 2016). As a result, the overall variance of the measured methylation values across all the CpG sites in the studied population can be described as:

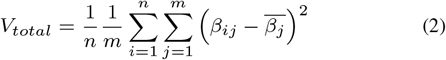

Where *V_total_* represents the total variance of the studied samples, *n* is the total number of all samples, m is the total number of studied CpGs, *β_ij_* represents the methylation beta value of the *j^th^* CpG in the *i^th^* sample, 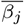 represents the mean methylation beta value of the *j^th^* CpG across all samples. Theoretically, we can then split the overall variance into the following two parts:

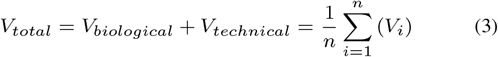

The first part *V_biological_* represents variance caused by meaningful biological reasons, such as cell types, age, gender, health status and other reasonable factors. The second part *V_technical_* represents variance resulting from technical issues, such as batch effect, random fluctuation and other unknown issues.

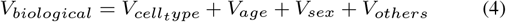

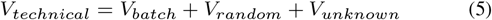

Sex is one of the major biological factors which influences the methylation status of many autosomal CpGs, as a result, hundreds of autosomal CpGs have been reported showing significant different methylation levels between sexes (McCarthy *et al.*, 2014; Yousefi *et al.*, 2015; Grant *et al.*, 2021). The fraction of variances which are explained by sex can be deduced as follows:

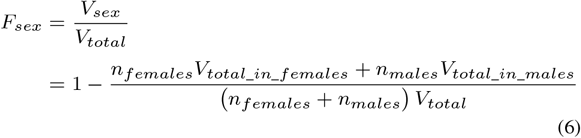

Ideally, a good normalisation method should be able to not only greatly reduce the variances that are resulted from technical issues (*V_technical_*), but also need to keep variances which have meaningful biological reasons (*V_biological_*). This means, after the normalisation process, the overall variance should be reduced significantly while the sex explained fraction of variance should be increased. In this paper, to study the potential sex bias introduced by the mix normalisation method dasen, we compared the mean variance and the fraction of sex explained variances of the methylation values of CpGs after no normalisation (raw beta values), dasen normalisation and interpolatedXY adjusted dasen normalisation within the three chromosome groups (i.e. autosomes, X chromosomes and Y chromosomes).

### 2.6 Artifactual sex differences

If the conventional mixed normalisation approaches do introduce systematic artificial sex biases into the autosomal CpGs, then some autosomal CpGs could be falsely sex-associated. Epigenome-wide association studies (EWAS) are commonly used to systematically assess the association between DNA methylation levels at genetic loci across the genome and a phenotype of interest. In this study, we apply EWAS to identify sex-associated CpG sites and then compare the EWAS results resulted from different preprocess approaches.

To perform EWASs for sex, the *champ.dmp* function in champ package (Tian *et al.*, 2017), which utilises linear regression and F-test to identify differentially methylated positions is applied in this study to identify sex-associated CpGs. After Bonferroni multiple comparison correction, those CpG sites with *p*-value less than 0.05 were selected as significantly sex-associate. For simplicity and better comparison, we do not include age, cell type proportions and other covariates within the EWASs.

### 2.7 Comparison of the original funnorm and the interpolatedXY adjusted funnorm

Funnorm is reported to be suitable for normalising methylation data with substantial global differences. The main difference between the original funnorm and the proposed interpolatedXY adjusted funnorm is how to normalise the methylation values of sex chromosome linked CpGs. The original funnorm is designed to normalise X chromosomes separately and differently with Y chromosomes, as well as processes female samples and male samples separately. In contrast, the interpolatedXY adjusted funnorm does not require prior sex annotations and process both genders equally, which generates the corrected values of sex chromosome linked CpGs by interpolation on the normalised values of autosomal CpGs.

To compare the normalisation effects on sex chromosome data between the original funnorm and the adjusted funnorm, we studied both the density distributions and the variances of the methylation values of CpG sites after no normalisation (raw beta values), funnorm normalisation and adjusted funnorm normalisation within three chromosome groups (i.e. autosomes, X chromosomes and Y chromosomes) in two 450k datasets. The first dataset (dataset one) includes 12 male samples and 4 female samples, while the second dataset (dataset two) contains 23 male samples and 25 female samples.

**Fig. 2:**
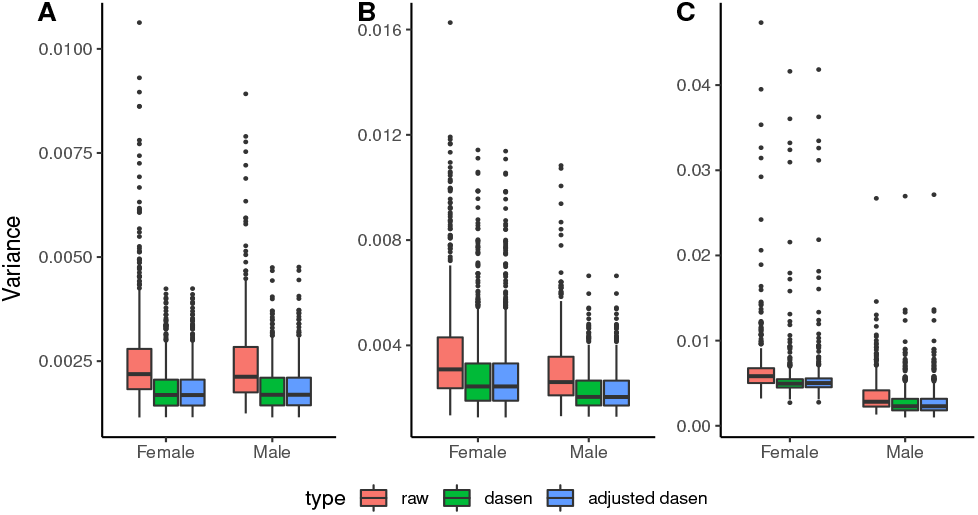
Variance comparisons in the UKHLS dataset. Boxplots comparing the variance of methylation beta values with three different pre-processing methods (i.e. no normalisation, dasen normalisation and adjusted dasen normalisation) in autosomes (A), X chromosomes (B) and Y chromosomes (C). Females and males are dealt with separately.

## 3 Results

### 3.1 Estimation using the interpolation approach

We first investigated the performance of the interpolation approach employed by the interpolatedXY adjusted funnorm method. The deviations from the inferred values by the interpolation approach to their corresponding reference values are measured by RMSE. As it can be seen from Figure 1, the resulting RMSEs are all very small, especially for those in both X chromosomes and male Y chromosomes: the mean RMSE of X chromosome linked CpGs is 1.15e-05 (sd=8.7e-06) in females and is 1.11e-05 (sd=4.8e-06) in male samples, while the mean RMSE of estimations for male Y chromosomes is 6.61e-06 (sd=3.2e-06). Though the RMSEs of Y chromosome linked CpGs in females are slightly higher (mean=8.98e-04, sd=6.0e-04), they are still very subtle. With the knowledge that females do not carry Y chromosomes, and those observed signal intensities result from background noises and non-specific hybridization, there is no need to look much into the methylation values of female Y chromosomes. In the same way, we could observe similar performances of the interpolation approach employed by the interpolatedXY adjusted dasen method (Supplementary Figure 1).

**Fig. 1:**
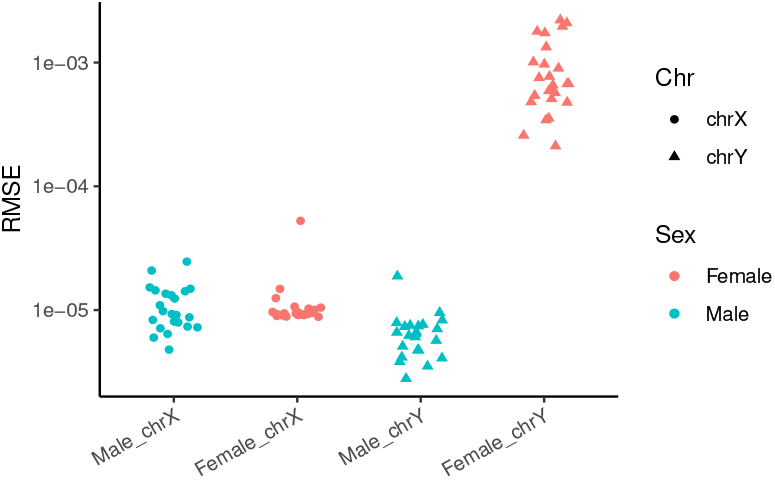
Difference between interpolated values and expected values within the adjusted funnorm. RMSEs are grouped in four categories: male X chromosomes, female X chromosomes, male Y chromosomes and female Y chromosomes. Female samples are in red colour and male samples are in blue colour. Dots represent X chromosomes, while triangles represent Y chromosomes.

In summary, the above results demonstrate the proposed interpolation approach provides accurate and robust estimations for the corrected values of sex chromosome linked CpGs.

### 3.2 Artificial sex biases are introduced into autosomal CpGs by the conventional mixed normalisation method

The first round of the UKHLS dataset (Gorrie-Stone et al., 2019) includes 1175 whole blood samples whose DNA methylation levels were measured using the EPIC array. After quality control, 685 female samples and 486 male samples were kept for this analysis. To study the normalisation effects, the variance of beta values with three different pre-processing methods (no-normalisation, dasen and interpolatedXY adjusted dasen) are compared within three different chromosome groups (i.e. autosomes, X chromosomes and Y chromosomes) separately. As shown in Figure 2, both dasen and adjusted dasen significantly (Wilcoxon signed-rank test, *p*-value less than 2.2e-16) reduce the variance in all three chromosome groups. For instance, the mean variance of autosomes in both sexes decreased from around 0.0025 in non-normalised beta values to about 0.0018 after either dasen or adjusted dasen normalisation. The beta values density plots also demonstrate that both dasen and adjusted dasen greatly reduce the distribution variation (e.g. Supplementary Figure 2). However, the difference in normalisation effects between dasen and adjusted dasen is not significant from the variance level.

Table 1 describes the sex explained fraction of variance between three methods in three chromosome categories. We can see that the sex explained variance in sex chromosomes by the three methods all exceeds 70%, while it accounts to only around 0.5% in autosomes. That is in line with our expectation, as sex is a dominant factor causing difference in methylation levels of sex chromosomes, while the majority of autosomal CpGs are not influenced by sex. Interestingly, the sex explained fraction of variance of raw beta values in autosomes is 0.34%, it rises to 0.45% after normalising by the adjusted dasen, indicating the adjusted dasen method retained the meaningful biological difference when reducing technical variances (Figure 2A). However, the sex explained variance is much higher (0.57%) by normalising with the original dasen, cthe an we conclude that the original dasen is better than the adjusted dasen to retain meaningful biological difference? On the contrary, these results indicate the original dasen has introduced artificial sex bias into to the normalised data. Combining the facts that only autosomal CpGs were included to compute the variance, and the difference in normalising the autosomal CpGs between the two methods is that the correction of autosomal CpGs is affected by the enrolling of sex chromosome data within the original dasen procedures, but not influenced within the adjusted dasen method. We can conclude that the observed higher fraction (sex explained fraction of variance in autosomes) with the original dasen normalisation is partly driven by the involvement of sex chromosome data, and this higher figure (i.e. than the adjusted dasen) indicates that technical sex biases have been introduced into to autosomal CpGs by the original dasen.

**Table 1.** The fraction of variance explained by sex in the UKHLS dataset with no normalisation (raw), dasen normalisation, interpolatedXY adjusted dasen normalisation and interpolatedXY adjusted funnorm normalisation.

| Fraction of variance explained by sex (%) | Raw | Dasen | Adjusted dasen | Adjusted funnorn |
| --- | --- | --- | --- | --- |
| Autosomes | 0.34 | **0.57** | 0.45 | **0.46** |
| X chromosome | 73.18 | 77.24 | **77.57** | 76.93 |
| Y chromosome | 85.34 | 87.64 | 87.50 | **88.82** |

### 3.3 Confirmation of the introduced sex biases

We performed EWASs of sex based on autosomal beta values of UKHLS samples with three different pre-processing: no normalisation, dasen normalisation and interpolatedXY adjusted dasen normalisation. The identified number of sex significant (Bonferroni *p*-value less than 0.05) differentially methylated positions (saDMPs) are shown in Figure 3.

**Fig. 3:**
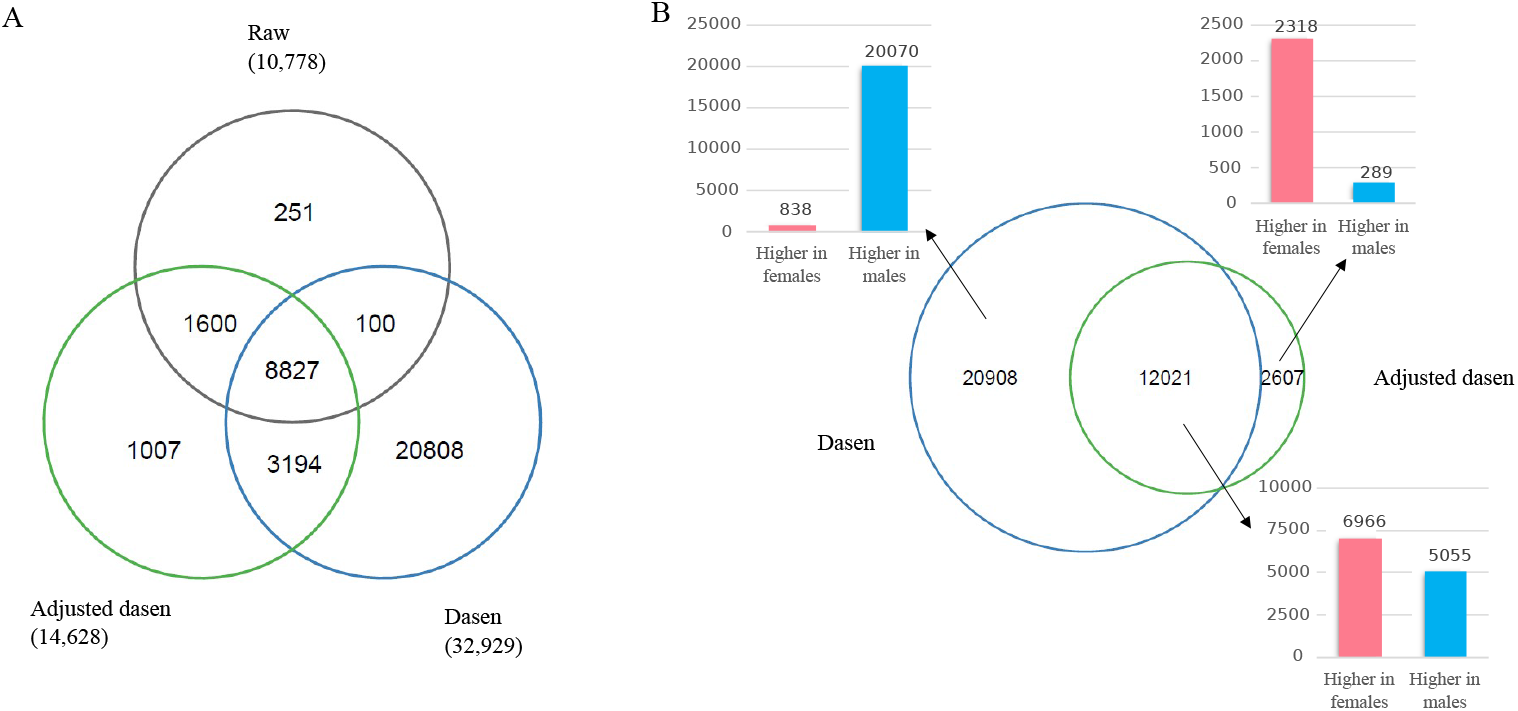
EWAS results of UKHLS dataset. A. The Venn diagram shows the number of unique and shared saDMPs between three approaches: no normalisation (raw), dasen normalisation and adjusted dasen normalisation. B. The Euler diagram describes the number of unique and shared saDMPs between dasen normalisation and adjusted dasen normalisation, with the three bar plots showing the number of CpGs which have higher methylation values in females (red) or males (blue) in three categories separately.

As illustrated in the Venn diagram (Figure 3A), there are 10,778 CpG sites been identified as saDMPs in the raw data, with 96.7% of them (10,427) also been captured after adjusted dasen normalisation. In addition, compared to raw data, the adjusted dasen approach enables the identification of another 4,201 saDMPs. Once again, these results demonstrate that while the adjusted dasen greatly reduces the variation of beta values (Figure 2A), it preserves the meaningful biological differences.

We found a total of 32,929 saDMPs after the original dasen normalization, which is more than three times the number with no normalisation or 2.25 times the number with adjusted dasen normalisation. Even so, 1,600 CpGs which are identified by both no normalisation and adjusted dasen normalization, are missed by the original dasen method. When comparing the dasen and adjusted dasen (Figure 3B), there are 12,021 saDMPs shared between the two methods. Interestingly, among the 20,908 dasen specific saDMPs, 96.0% of them (20,070) have higher methylation values in males than that in females. On the contrary, 2,318 out of the 2,607 adjusted dasen specific saDMPs (88.9%) show higher methylation values in females than males. Again, with the fact that the interpolatedXY adjusted dasen only differs from the original dasen by not enrolling sex chromosome data when normalising the autosomal data, the above results suggest the original dasen did introduce artificial sex biases into autosomal CpGs by making the methylation values of many CpGs slightly higher in male samples and lower in female samples. This explains why nearly all the dasen specific saDMPs have higher methylation values in male samples, and there are more than two thousand CpG sites which have higher methylation values in female samples that were identified as significant saDMPs by the adjusted dasen approach but missed by the original dasen.

### 3.4 InterpolatedXY adjusted funnorm provides better normalisation results for sex chromosome linked CpGs than the original funnorm

Since the original funnorm has two different designs to deal with different size datasets, we compared the normalisation effects between the original funnorm and the interpolatedXY adjusted funnorm in two datasets. The adjusted funnorm does not differ from the original funnorm in normalising the autosomal CpGs, so the corrected values of autosome data from the two methods are the same, we can thus observe identical results for autosomal CpGs by the two methods (Figure 4B and 4C, Table 2, Supplementary Figure 3B and 3C, Supplementary Table 2).

**Fig. 4:**
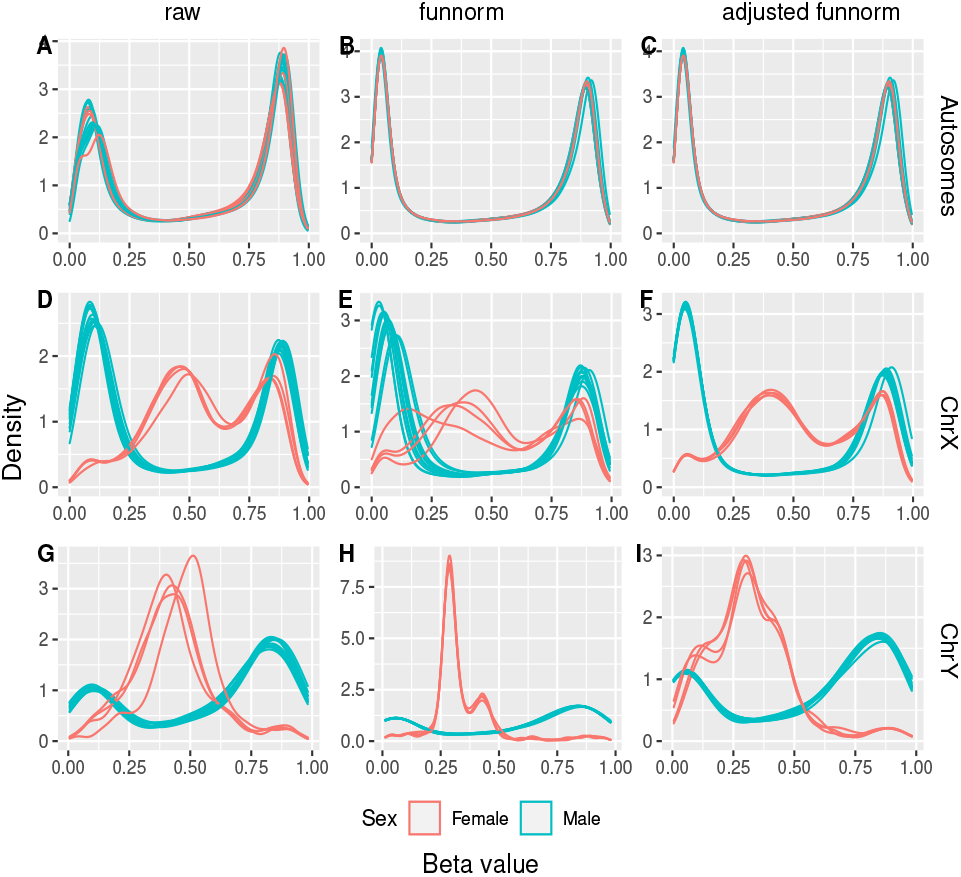
Comparisons in methylation beta value density distributions for dataset one. The three columns list results from raw data (left column), funnorm normalised data (middle column) and the adjusted funnorm normalised data (right column). The three rows show density distributions of autosomal CpGs (first row), X chromosome linked CpGs (second row) and Y chromosome linked CpGs (third row). Red lines represent females and blue lines represent males.

**Table 2.** The fraction of variance explained by sex in the dataset one (n=16) with no normalisation (raw), funnorm normalisation and interpolatedXY adjusted funnorm normalisation.

| Fraction of variance explained by sex (%) | Raw | Funnorm | Adjusted funnorm |
| --- | --- | --- | --- |
| Autosomes | 9.48 | 10.93 | 10.93 |
| X chromosome | 92.68 | **82.82** | 92.99 |
| Y chromosome | 91.48 | **97.09** | 93.89 |

For the X chromosome linked CpGs, when applied to small datasets, whose number of female samples or male samples is less than ten, such as dataset one, funnorm is designed to normalise female X chromosomes and male X chromosomes together by the functional normalisation. Compared to the non-normalised raw beta values, the density distributions of the corrected data generated by funnorm turn out to be much discordant in both female samples and male samples (e.g. Figure 4E). On the contrary, after the adjusted funnorm normalisation, the density distributions become more consistent in both sexes (Figure 4F). We can also observe the same trends from the bar plots in Figure 5B, the original funnorm greatly increases the variance in both sex groups, while the adjusted funnorm keeps the variance low. Furthermore, the sex explained fraction of variance reduced to 82.8% by the original funnorm, which is 92.7% in raw data and 93.0% after the adjusted funnorm normalisation (Table 2). Taken together, the above results indicate that the original funnorm is actually adding technical variation into the methylation data of X chromosomes for those small sample size datasets.

**Fig. 5:**
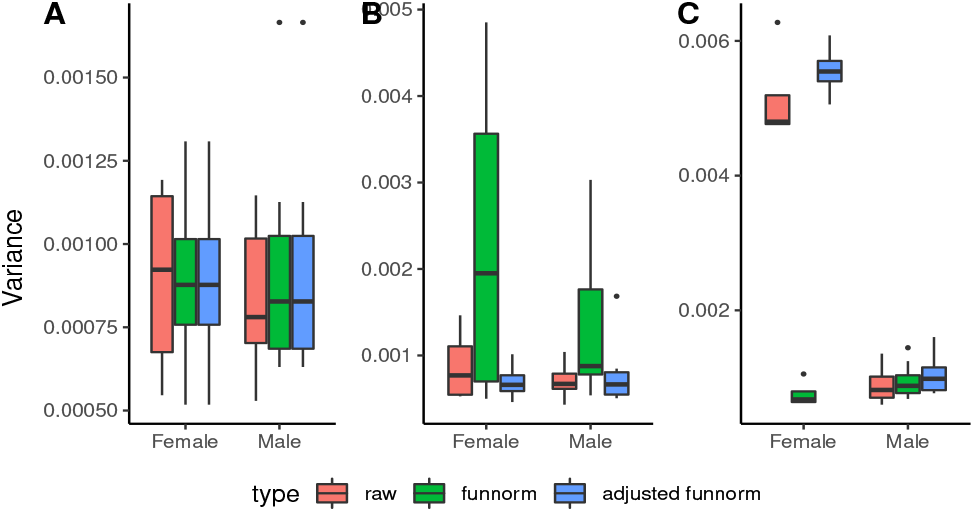
Variance comparisons in the dataset one. Boxplots comparing the variance of methylation beta values with three different pre-processing methods (i.e. no normalisation, dasen normalisation and adjusted dasen normalisation) in autosomes (A), X chromosomes (B) and Y chromosomes (C). Females and males are dealt with separately.

**Fig. 6:**
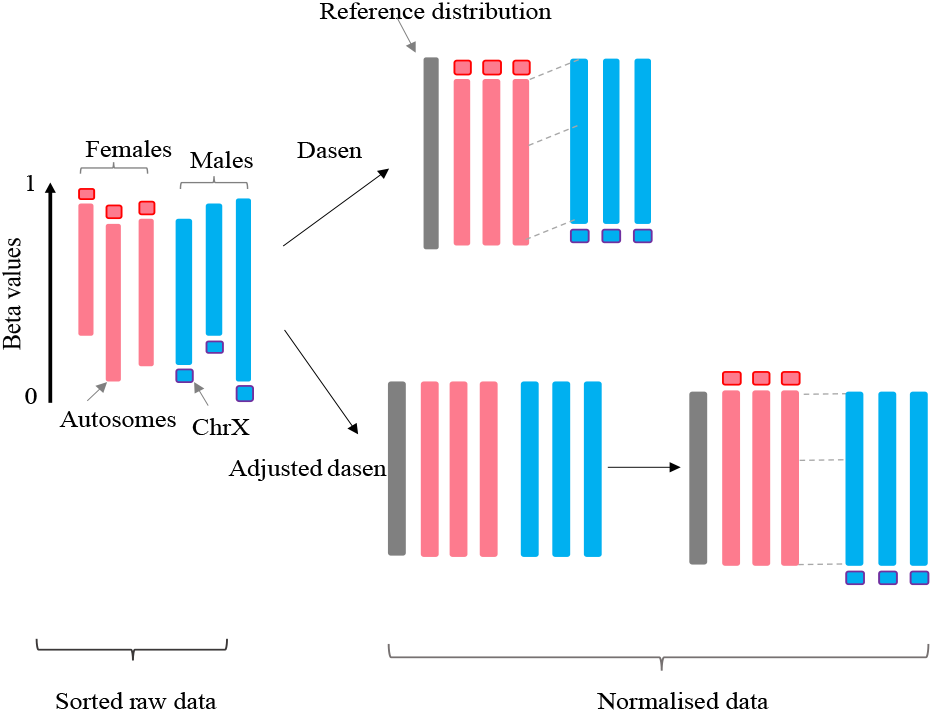
A simplified schematic diagram illustrates the difference in the normalisation process between the original dasen and the interpolatedXY adjusted dasen. The original dasen normalises autosomes and sex chromosomes together, the mean methylation values of most X chromosome linked CpGs in females are higher than nearly half of the autosomal CpGs, whereas the values of the corresponding locus in males are relatively very low, thus the quantile normalisation algorithm employed by dasen to make all studied samples fit into a same distribution creating a systematic shift for many autosomal CpGs in two sexes. The adjusted dasen manages to avoid such an issue by doing quantile normalisation in autosomes separately and independently with sex chromosomes, and infer the corrected values of sex chromosomes by interpolating on autosomes. Red denotes female sample and blue denotes male sample, the long bar represents sorted autosomal CpGs and the short bar represents sorted X chromosome linked CpGs.

When applied to larger datasets, such as in the case of dataset two, funnorm performs separate functional normalisations on female X chromosomes and male X chromosomes, with the underlying consideration that females and males have very different methylation patterns on X chromosomes. When comparing the normalisation effects between the original funnorm and the adjusted funnorm based on dataset two, we did not observe any significant differences in the methylation profiles of X chromosomes (Supplementary Figure 3, Supplementary Figure 4 and Supplementary Table 2).

For the Y chromosome linked CpGs, the original funnorm does not use the functional normalisation as it does on other chromosomes, such as autosomes. Instead, only quantile normalisation is employed by the original funnorm to normalise the Y chromosome data, and with female samples and male samples processed separately. This may explain why the sex explained variance within the original funnorm is much higher (i.e. 97.7%) than that in the raw data (i.e. 88.5%) and adjusted funnorm (i.e. 89.1%) (Supplementary Table 2). We can also observe similar trend from Table 2. These results suggest the separate normalisation strategy employed by the original funnorm will increase the difference between the two sex groups, and thus introduce artificial technical bias.

### 3.5 Comparison between the interpolatedXY adjusted funnorm and interpolatedXY adjusted dasen

We have demonstrated that the fraction of variance explained by sex is very useful to measure the normalisation effects for different methods and have also shown that the adjusted the dasen and the adjusted funnorm are both superior than their original versions. Then we compared their normalisation effects on a large healthy population: the UKHLS dataset (n=1171). The results are shown in Table 1, the first obvious observation is that both the adjusted dasen and the adjusted funnorm clearly increased the fraction of variance explained by sex in all chromosome groups (i.e. autosomes, X chromosome and Y chromosome) than the raw data, demonstrating that the use of either normalisation method is beneficial and worthwhile. As compared to the two adjusted normalisation methods, we can see their effects are comparable in the studied dataset (Table 1): the adjusted funnorm marginally outperforms the adjusted dasen in normalising the autosome data (0.46% vs. 0.45%) and Y chromosome data (88.82% vs. 87.5%), while the adjusted dasen is slightly better in normalising the X chromosome data (77.57% vs. 76.93%).

## 4 Discussion

We have described a two-step sex-unbiased data normalisation strategy for normalising DNA methylation microarray samples, which can be applied into almost all quantile-based normalisation methods, such as dasen and funnorm. By this strategy, the autosomal CpGs are normalised independently and separately from the sex chromosome CpGs, while the corrected values of sex chromosomes CpGs are estimated as the weighted average of the corrected methylation values of their nearest neighbour atusosomal CpGs.

The two steps are necessary. Since the average methylation levels of CpGs on X chromosome in females are very different from that in males, normalising them together with the autosomal CpGs, especially by the quantile-based methods, will introduce technical biases for both autosomes and sex chromosomes. By comparing the normalisation effects of the original dasen and the interpolatedXY adjusted dasen, we confirmed that the technical sex biases were introduced into the autosomal CpGs by the mix normalisation approach (original dasen)–with the sex explained fraction of variance in autosomes rising to 0.57% from 0.44% in the adjusted dasen normalised data. We further propose a rational explanation for this: within the quantile normalisation steps in dasen, there are procedures to sort and return ranks for all the probes, as the mean methylation values of the most X chromosome linked CpGs in females are higher than nearly half of the autosomal CpGs, whereas the methylation values of the corresponding positions in males are relatively low, thus the quantile normalisation algorithm used to make all studied samples fit into a same distribution creating a systematic negative shift for many autosomal CpGs (their methylation values are lower than most X chromosome linked CpGs) in females and a systematic positive shift for those CpGs in males. As a result of this, when we perform EWAS to look for autosomal sex-associated CpGs, the original dasen approach identified more than two times the number as identified by the adjusted dasen or non-normalised data. Moreover, 96.0% of the dasen specific saDMPs show higher methylation values in male samples than in female samples, by contrast, the majority of the 2,607 CpGs missed by the original dasen but identified by the adjusted dasen have higher methylation values in female samples than male samples.

Estimation of the corrected values for sex chromosomes CpGs by looking at their nearest neighbours on autosomes is made both possible and reliable by the fact that DNA methylation microarrays simultaneously measure over half a million CpG sites across the genome, and only a relatively small portion (i.e. 2.3% in EPIC and 2.4% in 450K) is mapped on the sex chromosomes. Here in this study, we have demonstrated that the linear interpolation approach provides both accurate and robust estimations for the sex chromosome data, with the mean RMSE less than 1.2e-5.

Funnorm is favoured for normalising methylation data with substantial global differences, such as cancer samples (Fortin *et al.*, 2014). With the consideration that females and males have distinct methylation patterns for sex chromosomes, funnorm has very explicit rules to normalise X chromosomes and Y chromosomes differently. Within the functional normalisation in funnorm, there is a regression step to infer the explainable technical variants based on control probes. The authors may have considered the regression models would be less accurate in the circumstance of only few samples, so funnorm is designed to perform functional normalisations on female X chromosomes and male X chromosomes together when the number of either female samples or male samples is less than ten. Our results in Section 3.4 have clearly shown that such a mix normalisation approach is destructive to the methylation profiles of X chromosomes in both females and males. Though to do functional normalisation on females and males separately is a way to avoid such an issue, it may also introduce potential systematic technical bias between the two separate groups.

For the Y chromosome linked CpGs, the original funnorm does not actually perform the functional normalisation as it does on other chromosomes, instead it performs only quantile normalisations on Y chromosomes, and processes female samples and male samples separately. As the proposed interpolatedXY adjusted funnorm could provide near-perfect estimations for corrected values generated by functional normalisation, it could be particularly useful for studies that focus on sex chromosomes DNA methylation data, especially when the methylation difference between the studied groups that are known to be very different. Moreover, by the adjusted funnorm method, the corrected values of sex chromosome linked CpGs are produced by linear interpolating on the distribution of autosomal CpGs, so in theory they are more comparable with the autosomal CpGs.

In this paper, we not only present a novel two-step strategy to unbiasedly normalise DNA methylation microarray samples, but also provide a useful concept-–the fraction of variance explained by sex, to quantitively measure the normalisation effect. Sex is an important biological factor that not only determines the methylation status of sex chromosomes, but also influences many autosomal CpGs. A good candidate normalisation method should not only be able to greatly reduce the technical variation between samples, but also should preserve the meaningful variation that has biological reasons (e.g. sex). Even though quantile normalisation has been widely employed by several DNA methylation normalisation methods, such as SWAN (Maksimovic *et al.*, 2012), dasen (Pidsley *et al.*, 2013) and funnorm (Fortin *et al.*, 2014). There are still concerns about whether the use of between-array normalisation methods could bring enough benefits to counterbalance the potential impairment of data quality (Dedeurwaerder *et al.*, 2013). Here, in this study, we demonstrated that the interpolatedXY adjusted dasen and the the interpolatedXY adjusted funnorm are two good normalisation method candidates, they are able to not only greatly reduce technical variation but also retain the meaningful biological difference, which will be very useful for large cohort EWAS projects.

We believe that the proposed novel two-step strategy may have wider application outside of DNA methylation microarrays and could even be applied in more broader technologies such as RNA-Seq.

## 5 Conclusion

The proposed two-step strategy of interpolatedXY provides an excellent solution to normalise autosomal data and sex chromosome data without bias. The two steps are necessary and reliable. With the introducing of the interpolatedXY, the adjusted dasen and the adjusted funnorm both show superior performance than their original versions, and the normalised data are much better than the non-normalised raw data to highlight the meaningful biological difference.

## Acknowledgements

The authors acknowledge the use of the High-Performance Computing Facility (Ceres) and its associated support services at the University of Essex in the completion of this work.

## Funding

The UK Household Longitudinal Study is led by the Institute for Social and Economic Research at the University of Essex and funded by the Economic and Social Research Council [ES/M008592/1]. LCS is funded by the Medical Research Council [MR/R005176/1]. KDM is funded by the Economic and Social Research Council [ES/M010236/1] and the Engineering and Physical Sciences Research Council [EP/P017487/1, EP/R02572X/1, EP/V000462/1]. XZ is funded by the Engineering and Physical Sciences Research Council [EP/V034111/1]. YW is funded by the University of Essex.

## Conflict of Interest

none declared.

## Notes

### Competing Interest Statement

The authors have declared no competing interest.

