## Supplemental Figures and Tables for "interpolatedXY: a two-step strategy to normalise DNA methylation microarray data avoiding sex bias"

### Supplementary table and figures

**Supplementary Table 1:** Lists of sample ID used in dataset one and dataset two.

| Sample | Sex | Dataset one(n=16) | Dataset two(n=48) |
|---|---|---|---|
| GSM4230892 | Female | TRUE | TRUE |
| GSM4230891 | Male | TRUE | TRUE |
| GSM4230890 | Male | TRUE | TRUE |
| GSM4230889 | Female | TRUE | TRUE |
| GSM4230888 | Female | TRUE | TRUE |
| GSM4230887 | Male | TRUE | TRUE |
| GSM4230886 | Male | TRUE | TRUE |
| GSM4230885 | Female | TRUE | TRUE |
| GSM4230884 | Male | TRUE | TRUE |
| GSM4230883 | Male | TRUE | TRUE |
| GSM4230882 | Male | TRUE | TRUE |
| GSM4230881 | Male | TRUE | TRUE |
| GSM4230880 | Male | TRUE | TRUE |
| GSM4230879 | Male | TRUE | TRUE |
| GSM4230878 | Male | TRUE | TRUE |
| GSM4230877 | Male | TRUE | TRUE |
| GSM4230876 | Male | FALSE | TRUE |
| GSM4230875 | Male | FALSE | TRUE |
| GSM4230874 | Female | FALSE | TRUE |
| GSM4230873 | Female | FALSE | TRUE |
| GSM4230872 | Female | FALSE | TRUE |
| GSM4230871 | Female | FALSE | TRUE |
| GSM4230870 | Female | FALSE | TRUE |
| GSM4230869 | Female | FALSE | TRUE |
| GSM4230868 | Female | FALSE | TRUE |
| GSM4230867 | Female | FALSE | TRUE |
| GSM4230866 | Female | FALSE | TRUE |
| GSM4230865 | Female | FALSE | TRUE |
| GSM4230864 | Female | FALSE | TRUE |
| GSM4230863 | Female | FALSE | TRUE |
| GSM4230862 | Female | FALSE | TRUE |
| GSM4230861 | Female | FALSE | TRUE |
| GSM4230860 | Female | FALSE | TRUE |
| GSM4230859 | Female | FALSE | TRUE |
| GSM4230858 | Female | FALSE | TRUE |
| GSM4230857 | Female | FALSE | TRUE |
| GSM4230856 | Female | FALSE | TRUE |
| GSM4230855 | Female | FALSE | TRUE |
| GSM4230854 | Female | FALSE | TRUE |
| GSM4230853 | Male | FALSE | TRUE |
| GSM4230852 | Male | FALSE | TRUE |
| GSM4230851 | Male | FALSE | TRUE |
| GSM4230850 | Male | FALSE | TRUE |
| GSM4230845 | Male | FALSE | TRUE |
| GSM4230844 | Male | FALSE | TRUE |
| GSM4230843 | Male | FALSE | TRUE |
| GSM4230842 | Male | FALSE | TRUE |
| GSM4230841 | Male | FALSE | TRUE |

**Supplementary Table 2:** The fraction of variance explained by sex in the dataset two (n=48) with no normalisation, funnorm normalisation and interpolatedXY adjusted funnorm normalisation.

| Fraction of variance explained by sex (%) | raw | funnorm | interpolatedXY adjusted funnorm |
|---|---|---|---|
| Autosomes | 6.39 | 7.05 | 7.05 |
| X chromosome | 93.33 | **93.33** | 93.21 |
| Y chromosome | 97.75 | **97.75** | 89.12 |

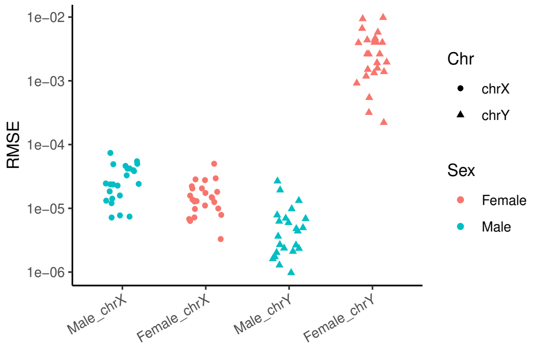


**Supplementary Figure 1: Difference between interpolated values and expected values within the adjusted dasen.** RMSEs are grouped in four categories: male X chromosomes, female X chromosomes, male Y chromosomes and female Y chromosomes. Dots represent X chromosomes and triangles represent Y chromosomes.

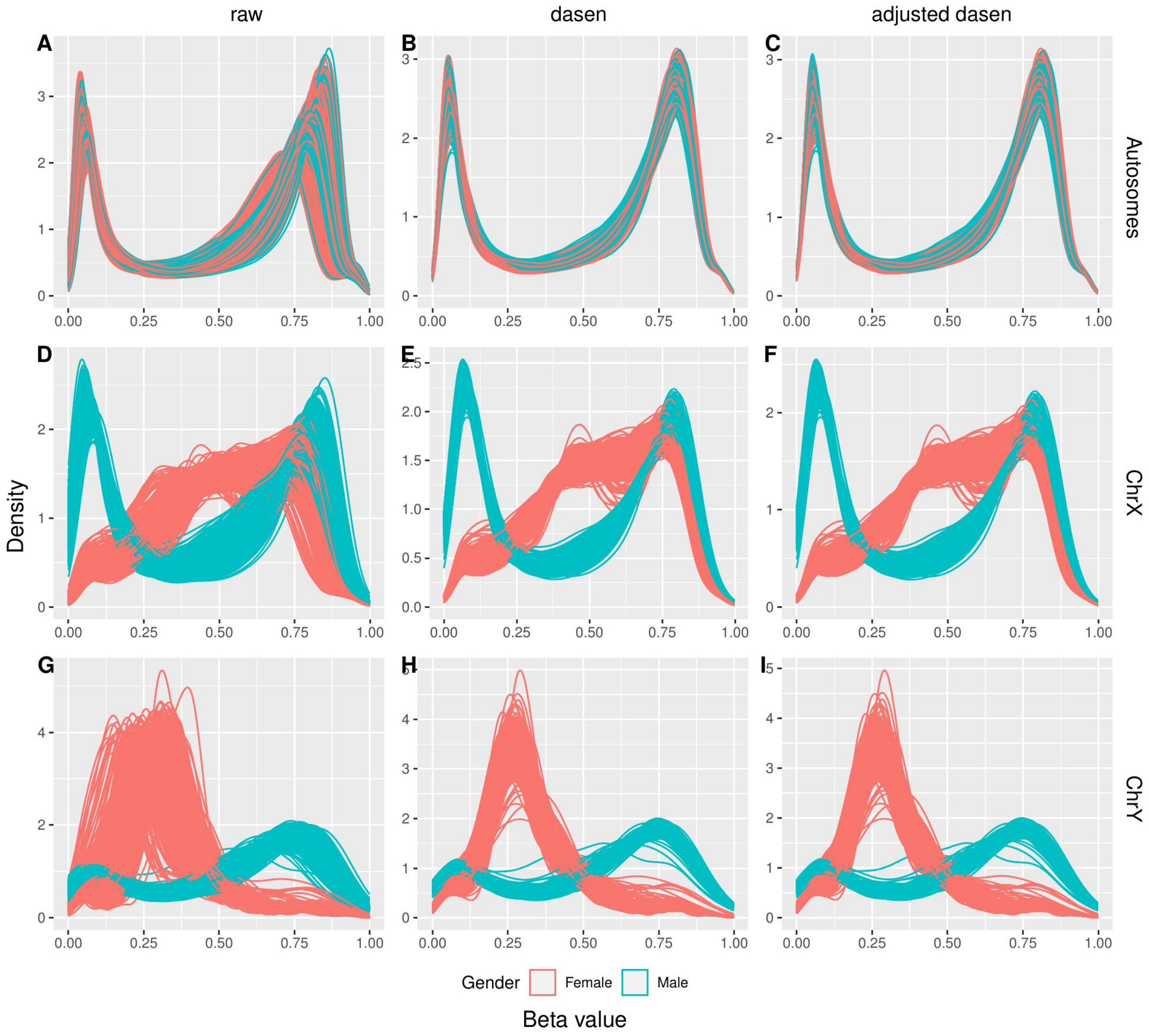


**Supplementary Figure 2: Comparisons in methylation beta value density distributions for UKHLS dataset.** The three columns illustrate results from raw data (left column), funnorm normalised data (middle column) and the adjusted funnorm normalised data (right column). The three rows show density distributions of autosomal CpGs (first row), X chromosome linked CpGs (second row) and Y chromosome linked CpGs (third row). Red lines represent females and blue lines represent males.

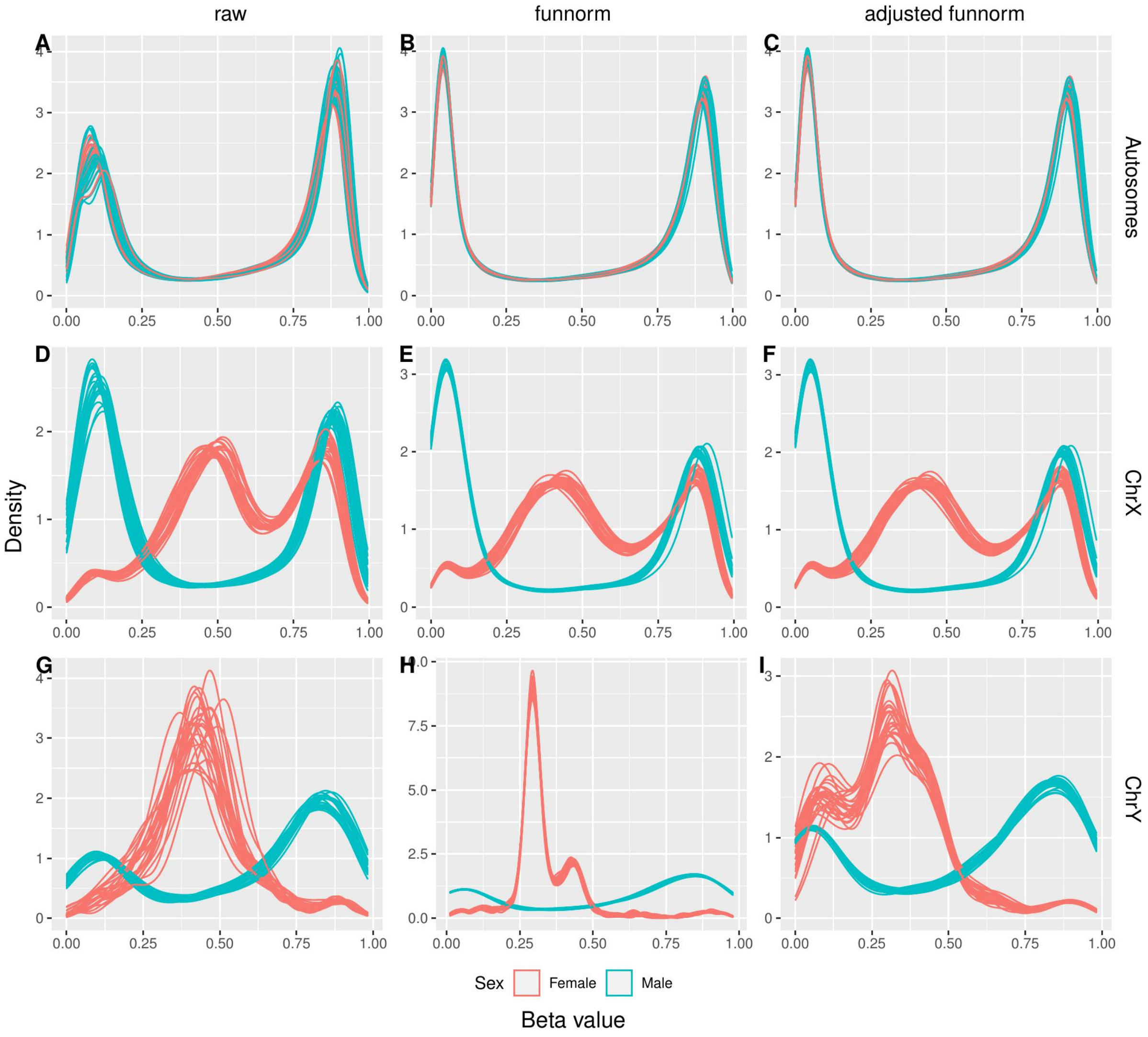


**Supplementary Figure 3: Comparisons in methylation beta value density distributions for dataset two.** The three columns illustrate results from raw data (left column), normalised data (middle column) and the adjusted funnorm normalised data (right column). The three rows show density distributions of autosomal CpGs (first row), X chromosome linked CpGs (second row) and Y chromosome linked CpGs (third row). Red lines represent females and blue lines represent males.

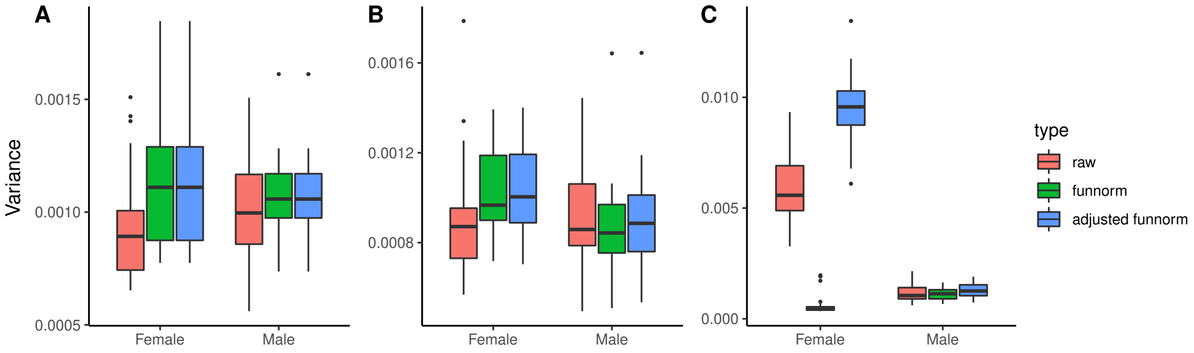


**Supplementary Figure 4: Variance comparisons in the dataset two.** Boxplots comparing the variance of methylation beta values with three different pre-processing methods (i.e. no normalisation, dasen normalisation and adjusted dasen normalisation) in autosomes (A), X chromosomes (B) and Y chromosomes (C). Females and males are dealt with separately.
